# Cross-species transmission and differential fate of an endogenous retrovirus in three mammal lineages

**DOI:** 10.1101/024190

**Authors:** Xiaoyu Zhuo, Cédric Feschotte

## Abstract

Endogenous retroviruses (ERVs) arise from retroviruses chromosomally integrated in the host germline. ERVs are common in vertebrate genomes and provide a valuable fossil record of past retroviral infections to investigate the biology and evolution of retroviruses over a deep time scale, including cross-species transmission events. Here we took advantage of a catalog of ERVs we recently produced for the bat *Myotis lucifugus* to seek evidence for infiltration of these retroviruses in other mammalian species (>100) currently represented in the genome sequence database. We provide multiple lines of evidence for the cross-ordinal transmission of a gammaretrovirus endogenized independently in the lineages of vespertilionid bats, felid cats and pangolin ∼13-25 million years ago. Following its initial introduction, the ERV amplified extensively in parallel in both bat and cat lineages, generating hundreds of species-specific insertions throughout evolution. However, despite being derived from the same viral species, phylogenetic and selection analyses suggest that the ERV experienced different amplification dynamics in the two mammalian lineages. In the cat lineage, the ERV appears to have expanded primarily by retrotransposition of a single proviral progenitor that lost infectious capacity shortly after endogenization. In the bat lineage, the ERV followed a more complex path of germline invasion characterized by both retrotransposition and multiple infection events. The results also suggest that some of the bat ERVs have maintained infectious capacity for extended period of time and may be still infectious today. This study provides one of the most rigorously documented cases of cross-ordinal transmission of a mammalian retrovirus. It also illustrates how the same retrovirus species has transitioned multiple times from an infectious pathogen to a genomic parasite (i.e. retrotransposon), yet experiencing different invasion dynamics in different mammalian hosts.

## INTRODUCTION

Viral cross-species transmission (CST) represents a major threat to both human and animal populations. Most viral diseases of humans are zoonotic: they stem from CST of viruses from domestic or wild animals [1]. The explosion and development of human society, including modern transportation, over the last 100 years has exposed us to an increasing number of pathogens [2]. AIDS, which has caused more than 25 million death over the past ∼30 years (aids.gov), is one of the most notorious examples of a pandemic initiated by viral CST [3,4]. The pathogens causing AIDS (HIV-1 and HIV-2) are retroviruses, a family of RNA viruses that use reverse transcription to replicate their genome [5]. Other retroviral CST events have been documented within primates, felids and ruminants, suggesting that retroviral CST represents a continuous threat to human and animal health [6-10].

Retroviruses are unique amongst animal viruses in that chromosomal integration of so-called proviruses is an obligatory step in their replication cycle [5]. As a consequence, retroviral infection of germ cells or their progenitors result in proviruses that may be vertically inherited along with the host genome. Such inheritable proviruses are called endogenous retroviruses (ERVs). Under some circumstances, which are still poorly understood, ERVs can further propagate within the genome and spread in the population, resulting in the formation of large families of interspersed repeats in the host genome [11]. Despite the potentially deleterious consequences associated with the genomic propagation of ERVs, the process has been remarkably pervasive during mammalian evolution. Indeed every mammalian genome thus far examined harbor a great abundance and diversity of ERVs, which are mostly lineage-specific. For example, 8% of the human genome is composed of ERV sequences derived from a wide variety of retroviruses acquired at different time points during primate evolution [12-14]. Once integrated and endogenized, most ERVs appear to evolve at the host’s neutral mutation rate, which is much slower than the mutation rate of exogenous retroviruses (XRVs) [15]. Therefore ERVs provide a valuable fossil record of past retroviral infections and a unique opportunity to investigate retroviral evolution at a deep time scale, including CST events [16-20].

Many ancient CST events have been inferred by comparing ERV sequences across species [21-28]. Most of the well-documented cases of retroviral CST events involve closely related host species (e.g. from the same order). Indeed, it is thought that viral CST is often constrained by the evolutionary distance between donor and recipient species [19,20,29]. The observation that all retroviruses known to infect humans have been acquired from other primates is consistent with this notion [7]. However, retroviral CST events can also occur between distantly related species. For example, the cat RD114 gammaretrovirus is a recombinant containing an envelope domain mostly closely related to Baboon endogenous virus (BaEV), and is thought have been acquired by the domestic cat from an Old World monkey [30,31]. Also, the koala retrovirus (KoRV), which is currently spreading and undergoing endogenization in the wild, is very closely related to gibbon ape leukemia virus (GALV) and to ERVs found in Asian rodents, from which it was most likely acquired [32]. It has also been reported that reticuloendotheliosis virus (REV) was likely transmitted from mammals to birds [10]. Recent phylogenomics surveys of ERVs across a wide variety of vertebrate species suggested that CST between widely diverged species (i.e. from different orders or classes) may be more common than initially anticipated [19,20,33]. However, the evidence remains limited and more detailed case studies are needed to confirm this idea.

Bats (order Chiroptera) are increasingly regarded as exceptionally potent reservoirs of zoonotic viruses [34-39]. Indeed, a variety of bat species have been implicated in the spillover of diverse and highly pathogenic RNA viruses such as Rabies, Nipah, Hendra, SARS, Marburg, and Ebola viruses in the human population [40]. Very recently, one potential case of CST of an endogenous betaretrovirus involving phyllostomid bats, rodents and New World monkeys was reported [28]. We previously produced a comprehensive catalog of ERVs in the vespertilionid bat *Myotis lucifugus* [41] (referred to as MLERVs hereafter), documenting a rich and recent history of retroviral infections in this species lineage. Here, we have taken advantage of this resource to seek evidence of CST events implicating MLERVs. We identified an intriguing case of a gammaretrovirus that colonized independently the genomes of vespertilionid bats, felids and pangolin but followed a different fate and amplification dynamics in these lineages.

## RESULTS

### ERVs closely related to MLERV1 are present in species from three mammal orders

To detect possible CST events involving *M. lucifugus* ERVs, we used the sequence of the reverse transcriptase domain (RVT_1) (642 nt) from members of each of the 86 MLERV subfamilies previously identified [41] as queries in megaBLAST searches of all mammal genomes deposited in the NCBI whole genome shotgun (WGS) database as of February 2015 (107 mammal species). Excluding hits to *M. lucifugus*, the most significant hits (>80% nucleotide identity over the entire domain; e-value < 10^-80^) were obtained with a query representing the MLERV1 family [41] against the genome assemblies of the domestic cat (*Felis catus*) [42], Amur tiger (*Panthera tigris*) [43] and Chinese pangolin (*Manis pentadactyla*). In addition, and less surprisingly, many highly significant hits to MLERV1 were also obtained in the genomes of vespertilionid bat species closely related to *M. lucifugus* (Brandt’s myotis, *Myotis brandtii* [44]; David’s bat, *Myotis davidii* [45]; big brown bat, *Eptesicus fuscus*).

Further examination revealed that the hits in the feline genomes corresponded to an endogenous gammaretrovirus family initially described in the domestic cat. Two proviruses of this family were initially documented in cat as FERVmlu1 and FERVmlu2 [46]. In 2011, this ERV family was also reported in Repbase [47] as ERV1-1_Fca. In a more recent and more systematic inventory of ERVs in the cat genome [48], this family was designated as FcERV_γ6, a nomenclature we will adopt hereafter. Most recently, this family was identified as part of “lineage VII” by Mata et al. [33] who also reported the presence of closely related gammaretroviral elements in several wildcat species, including jaguar, puma, jaguarundi and tiger. To our knowledge, the related elements in the pangolin have not been previously characterized elsewhere. Hereafter we refer to this novel ERV family as MPERV1 for *Manis pentadactyla* ERV1 and deposited its consensus sequence in Repbase. For simplicity, we refer to all the elements detected in vespertilionid bats as MLERV1 and all the elements in different felids as FcERV_γ6.

To determine the ERV copy number in each species, we used the LTR sequences to mask their corresponding genome assembly using the Repeatmasker program and parsed the positional output to infer the number of putative full-length proviruses (i.e. containing two LTRs) and solitary (solo) LTR (see Methods). The results of this analysis (Table 1) show that each species harbors a relatively small number of full-length proviruses (2-50) but often numerous solo LTRs (up to 1600+ in *M. lucifugus*). It should be noted that the vast majority of proviruses we inferred to be full-length (based on the occurrence of a pair of LTRs within 10 kb) contain sequencing/assembly gaps. Thus we cannot ascertain whether they contain all the coding domains of a complete provirus.

**Table 1:** Copy number of MLERV1 related proviruses in different species.

|  | Tiger | Cat | *E. fuscus* | *M. davidii* | *M. brandtii* | *M. lucifugus* | pangolin |
| --- | --- | --- | --- | --- | --- | --- | --- |
| Full-length<br>ERV | 55 | 88 | 3 | 48 | 51 | 204 | 2 |
| Solo LTR | 675 | 744 | 67 | 1042 | 948 | 1638 | 27 |
| total | 730 | 832 | 70 | 1090 | 999 | 1842 | 29 |

### A retroviral CST event involving bat, cat and pangolin

To illustrate the exceptional level of sequence similarity among MLERV1, FcERV_γ6 and MPERV1, we generated nucleotide pairwise alignments of FcERV_γ6 and MLERV1 and of FcERV_γ6 and MPERV1 using the most closely related full-length proviruses from each family and performed a sliding window analysis of nucleotide identity across the two pairwise alignments (Fig. 1a). As a comparison, we performed the same analysis for proviral sequences representative of HIV-1 (Group M subtype B) and its closest relative from the chimpanzee SIVcpz [49,50]. The results show that the two representatives of the MLERV1 and FcERV_γ6 families and two representatives of FcERV_γ6 and MPERV1 are highly similar throughout their entire length, with an average level of nucleotide identity (∼85%) comparable to that between HIV-1 and SIVcpz (Fig. 1a). The most divergent segment corresponds to the predicted surface (SU) domain of the envelope protein (∼70% identity in the N-terminal region and with two large indels in the central region). Elevated divergence in the SU region is also apparent between the two lentiviruses, as previously documented [51], and is thought to reflect the rapid adaptation of retroviral envelope to diverged host cell receptors [52]. In summary, MLERV1, FcERV_γ6 and MPERV1 are just as closely related to each other as HIV-1 and SIVcpz, and thus these three elements and their relatives in the bat, cat and pangolin genomes can be considered as endogenous elements descended from the same retrovirus.

**Fig. 1. High sequence similarity and taxonomic distribution of MLERV1, FcERV_γ6 and MPERV1**

(a) Sliding window analysis of percent sequence identity along pairwise alignments of entire proviruses. (b) Taxonomic distribution of MLERV1, FcERV_γ6 and MPERV1. A schematic of the phylogenetic relationship of the 55 species from the clade “Scrotifera” currently represented in the NCBI whole genome sequence database, with human and mouse shown as outgroups. The 55 species fall within 6 mammal orders: Pholidota, Carnivora, Cetacea, Artiodactyla, Perissodactyla, Chiroptera. Some of the species are collapsed by order/family with the number of species for each clade indicated into parentheses. The three independent retroviral invasions of MLERV1, FcERV_γ6 and MPERV1 are depicted above each of the mammal lineages affected.

**Fig. 1.**
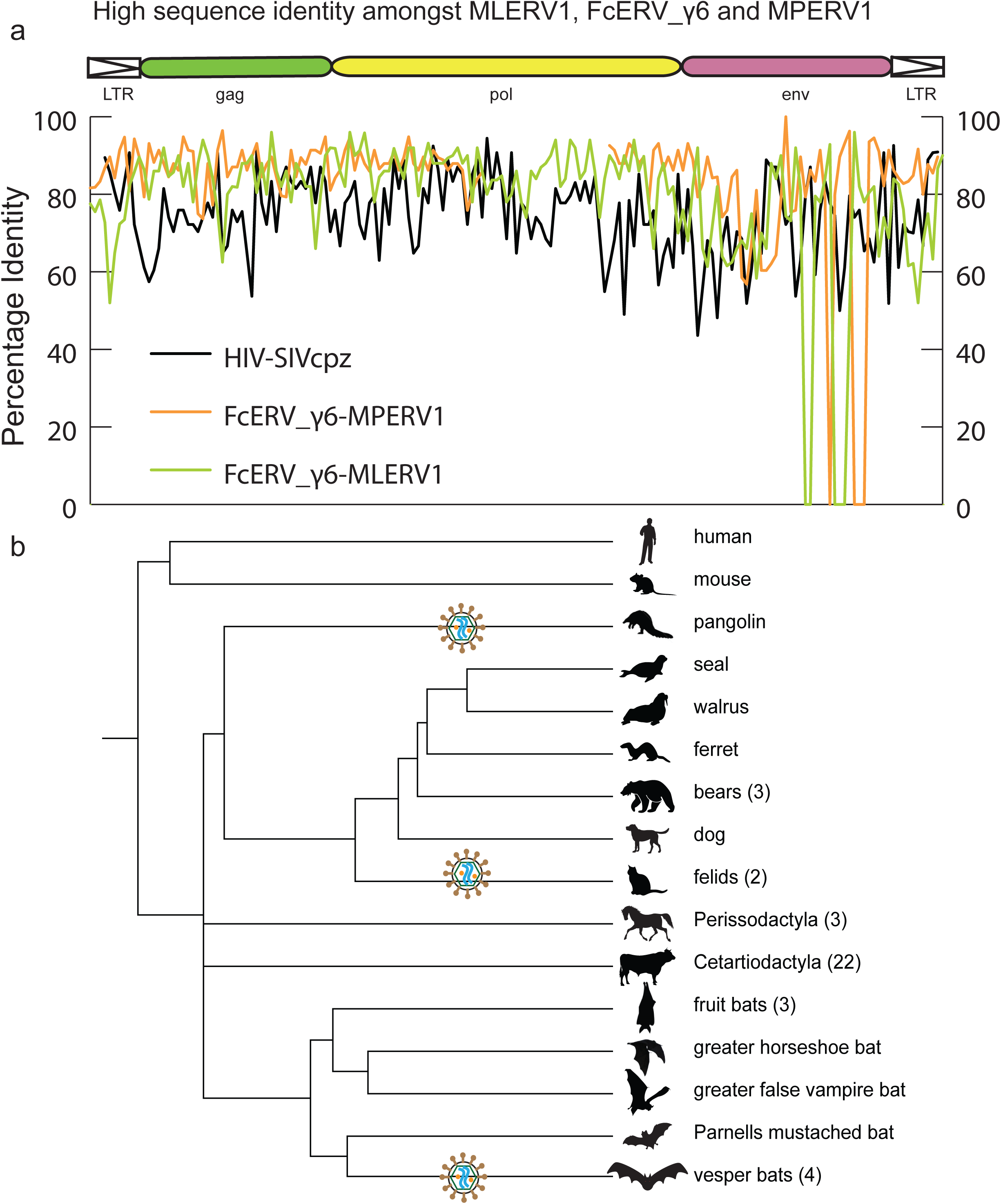
High sequence similarity and taxonomic distribution of MLERV1, FcERV_γ6 and MPERV1. a. Sliding window analysis of percent sequence identity along pairwise alignments of entire proviruses. b. Taxonomic distribution of MLERV1, FcERV_γ6 and MPERV1. A schematic of the phylogenetic relationship of the 55 species from the clade “Scrotifera” currently represented in the NCBI whole genome sequence database, with human and mouse shown as outgroups. The 55 species fall within 6 mammal orders: Pholidota, Carnivora, Cetacea, Artiodactyla, Perissodactyla, Chiroptera. Some of the species are collapsed by order/family with the number of species for each clade indicated into parentheses. The three independent retroviral invasions of MLERV1, FcERV_γ6 and MPERV1 are depicted above each of the mammal lineages affected.

The overall level of nucleotide similarity between MLERV1, FcERV_γ6 and MPERV1 is strongly incongruent with a scenario of vertical inheritance of an ancestral ERV present in the common ancestor of chiropterans, felids and pangolins, which dates back to ∼85 million year ago (MYA) [53,54]. Furthermore, we could not find any close relative of MLERV1 or FcERV_γ6 (no megaBLAST hit with sequence identity >80%) in the genome assemblies of species representative of other chiropteran (e.g. flying fox, Pteropodidae) or carnivore families (e.g. dog, Canidae; bear, Ursidae; ferret, Mustelidae; seal, Phocidae; walrus, Odobenidae). The Chinese pangolin genome is the only available representative of the order Pholidota, which is considered sister to Carnivora, and thus equally related to Perissodactyla (horse, rhino) and Cetartiodactyla (cow, pig, hippo, whales), all of which appear to lack related ERVs (Fig. 1b). Thus, the taxonomic distribution of MLERV1/FcERV_γ6 elements is extremely patchy, being detected in four vespertilionid bats, two feline species (cat and tiger), and one pangolin, but not in any of the numerous phylogenetically intermediate species represented in the NCBI WGS database (Fig. 1b). This taxonomic distribution suggests that the retrovirus that gave rise to MLERV1, FcERV_γ6 and MPERV1 underwent at least two CST events and was endogenized at least 3 times independently in the vespertilionid, felid, and pangolin lineages.

### Dating MLERV1/FcERV_γ6 insertions using comparative genomics

To gain further insights into the evolutionary history of these ERVs, we next sought to estimate when they first infiltrated their host genomes. Given that the likelihood of the same endogenous retrovirus to integrate at the same exact genomic location independently in different lineages is negligible, the presence of an element at orthologous position in different species can be interpreted as having inserted prior to their divergence time [55,56]. Conversely, since ERVs are not known to excise from the genome, the absence of an element in one species at a genomic location occupied by an ERV in another species strongly suggests that the ERV integrated after the split of the two species [57,58]. Such ‘empty’ site can be corroborated by the presence of a single copy of the host target sequence duplicated upon proviral integration (typically 4-bp target site duplication for gammaretroviruses). This cross-species presence/absence approach has been widely applied to date a variety of mobile element insertions, including ERVs [14,41,58,59].

We first examined the sharing of FcERV_γ6 elements between the cat and tiger, which diverged ∼10.8 MYA [60]. Out of a total of 1,419 putative full length proviruses and solo LTRs detected in the current whole genome assemblies of the two species, we were able to ascertain that 256 occupy orthologous positions, while 261 and 201 are specific to the cat and tiger lineages, respectively. None of these elements were detectable in other available carnivore genome assemblies (e.g. dog, panda, ferret, seal), while some of their flanking host sequences were readily detected (data not shown). These data indicate that FcERV_γ6 first invaded a felid ancestor sometime between ∼10.8 million years (MY) and ∼55 MYA and has continued to amplify to generate many insertions specific to the cat and tiger lineages (Fig. 2).

**Fig. 2. Distribution of MLERV1/FcERV_γ6 insertions in vesper bats and felids.**

The numbers of ERV insertions detected as orthologous or species-specific are shown as pies above each branch of the phylogeny of the vesper bats and felids examined. Different colors are used to illustrate FcERV_γ6 and the three different MLERV1 subfamilies.

**Fig. 2.**
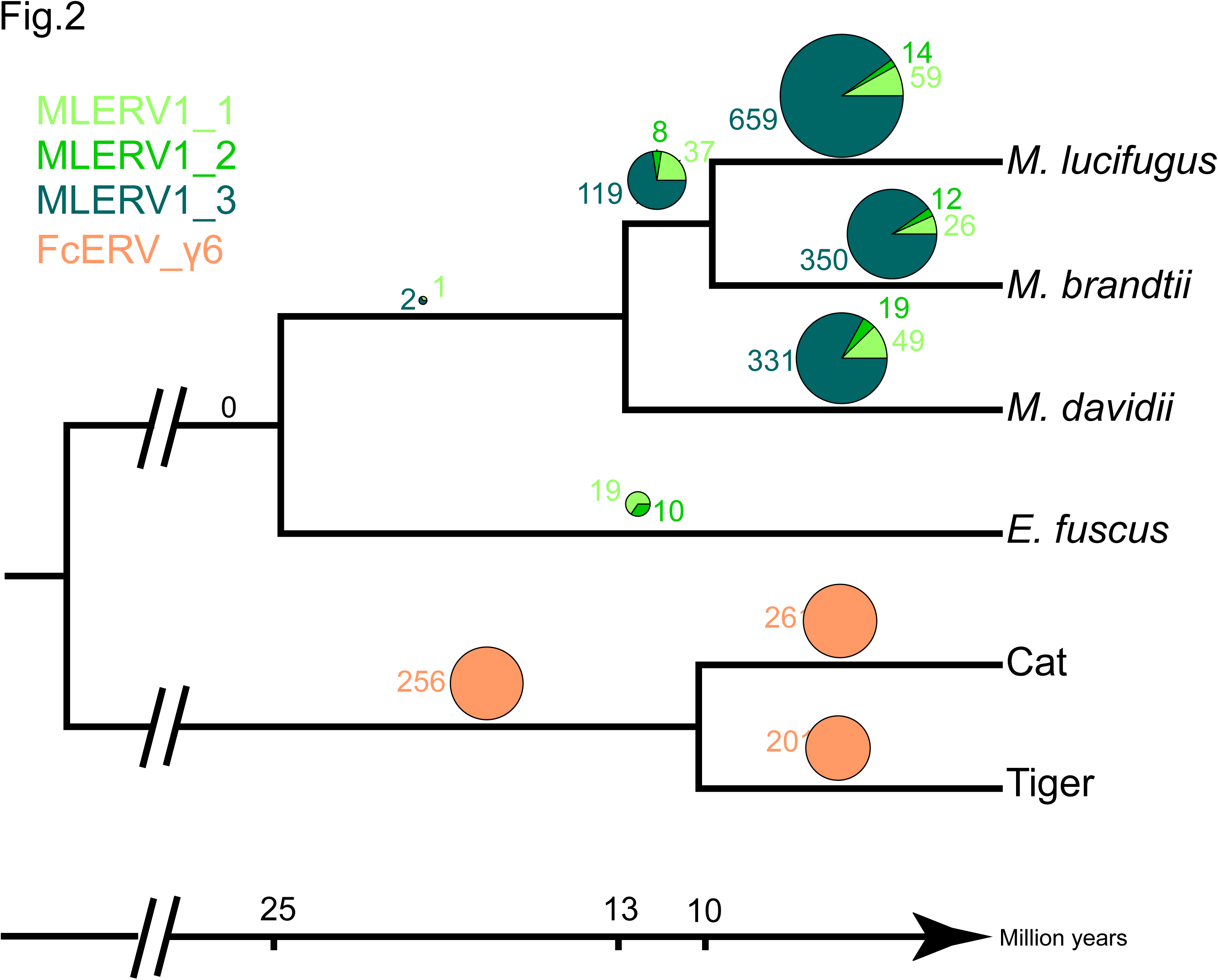
Distribution of MLERV1/FcERV_γ6 insertions in vesper bats and felids. The numbers of ERV insertions detected as orthologous or species-specific are shown as pies above each branch of the phylogeny of the vesper bats and felids examined. Different colors are used to illustrate FcERV_γ6 and the three different MLERV1 subfamilies.

Our previous phylogenetic analysis [41] has shown that the MLERV1 family of the little brown bat *M. lucifugus* can be divided into 3 subfamilies. Here we performed a systematic analysis of the presence/absence of MLERV1 elements (including solo LTRs) from the 3 subfamilies in the genome assemblies of three other vespertilionid bats currently available: Brandt’s myotis (*Myotis brandtii*), David’s bat (*Myotis davidii*) and the big brown bat (*Eptesicus fuscus*), which have been estimated to diverge from *M. lucifugus* ∼10 MYA, ∼13 MYA and ∼25 MYA, respectively [61-64]. The vast majority of MLERV1 elements and their close relatives were found to be species-specific (Fig. 2). Only 3 elements were present at orthologus loci across the 3 *Myotis* genomes (Fig. 2) and we could not find a single insertion shared between *E. fuscus* and any of the *Myotis*. Another interesting observation is that members of the MLERV1_3 subfamily, which contributes the vast majority (>80%) of MLERV1 elements in the 3 *Myotis* genomes, could not be identified at all in the *E. fuscus* genome. Indeed, all 29 elements detected in *E. fuscus* cluster with either one of the other two subfamilies (Fig. 4, and data not shown). Together these data suggest that the MLERV1 family expanded independently in the *Myotis* and *Eptesicus* lineages, but achieved a much higher copy number in the *Myotis* lineage due to the amplification of the MLERV1_3 subfamily, which has generated numerous species-specific insertions (Fig. 2).

**Fig. 3.**
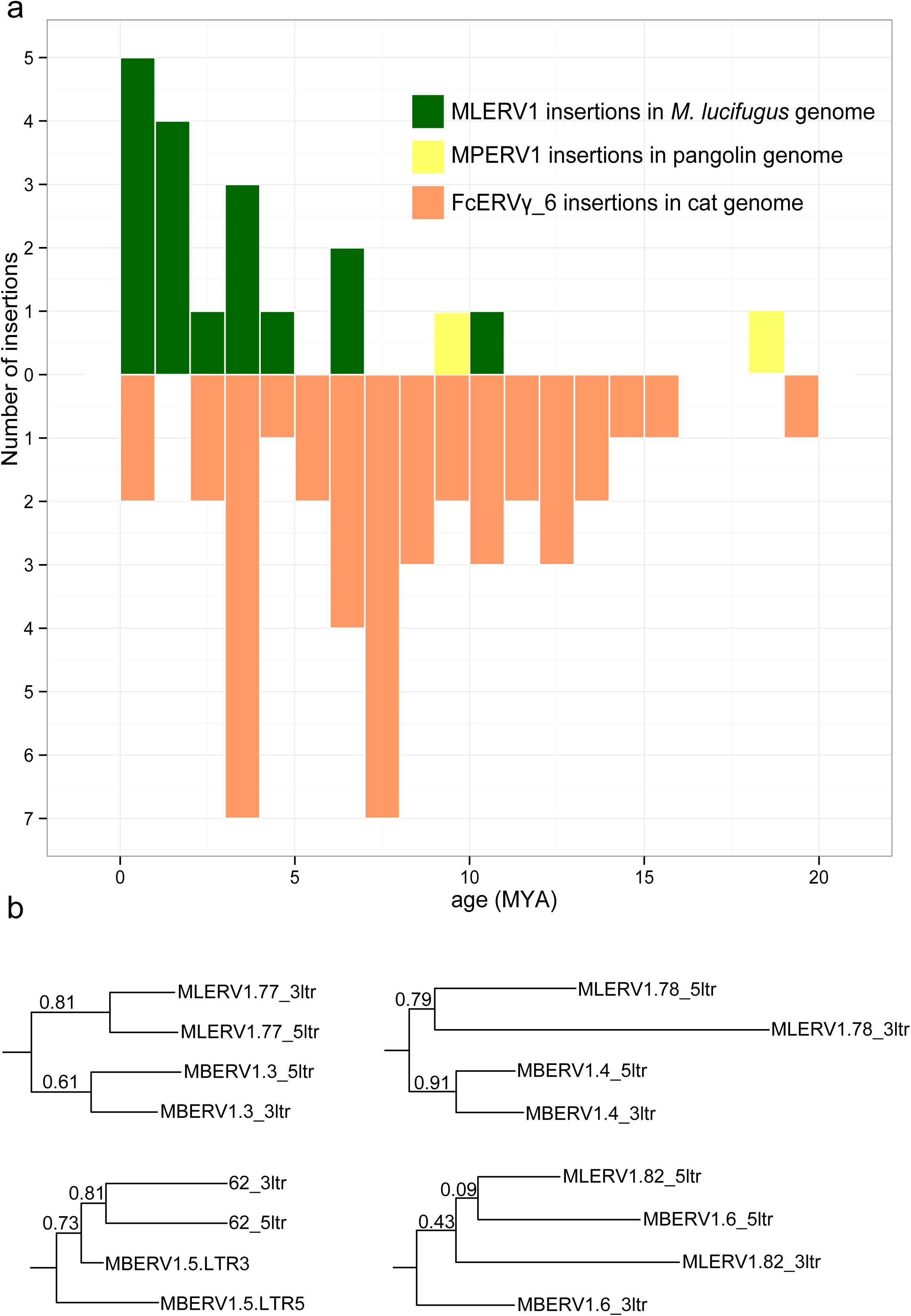
Dating individual proviral insertions based on LTR-LTR divergence. a. Age distribution of proviral insertions inferred from LTR-LTR divergence. The *y* axis shows the number of insertions for each age class binned in MY on the *x* axis. Each ERV family is shows as bars of different colors. b. Evidence of ‘gene’ conversion between 5’ and 3’ LTR of the same provirus. Four LTR trees are shown for four pairs of orthologous proviruses shared by *M. lucifugus* (MLERV) and *M. brandtii* (MBERV). Each maximum likelihood tree was built from a multiple alignment of the 5’ and 3’ LTRs from each provirus rooted with a non-orthologous LTR from *M. lucifugus* (not shown here to simplify the figure). The ML posterior probability is shown for each node. The fact that 5’ and 3’ LTR from the same provirus tend to group together rather than by species is indicative of gene conversion between the LTRs along the two species lineages following proviral insertion in their common ancestor.

**Fig. 4.**
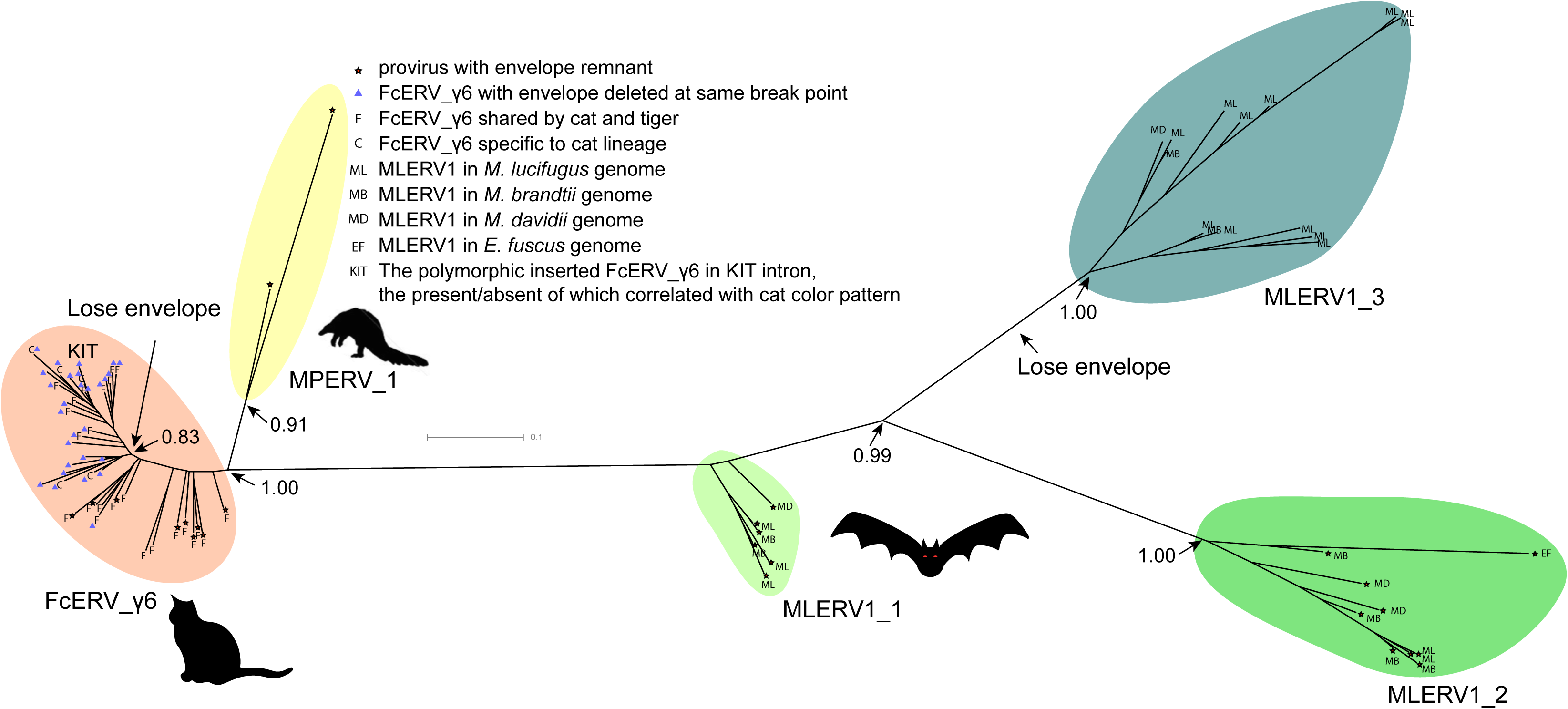
Phylogenetic analysis of MLERV1, FcERV_γ6, MPERV1 families. A maximum likelihood phylogenetic tree built from a multiple alignment of 3’ LTR sequences of 75 proviruses. The ML posterior probabilities supporting clades discussed in the text are shown. Information on the species origin as well as the presence/absence of envelope sequence is labeled at each node. Two independent losses of envelope by deletion are highlighted.

### Dating of individual provirus insertions using LTR-LTR divergence

Another widely applied method to date retroviral and other LTR-bearing retroelement insertions relies on the divergence of the 5’ and 3’ LTR of individual elements. This is because their retrotransposition mechanism results in two identical LTRs at the time of chromosomal integration. Given that most ERV LTR sequences are assumed to evolve neutrally once integrated in the host chromosome, the age of a provirus can be estimated based on LTR divergence by applying the host neutral substitution rate [48,58,65,66]. To eliminate the inflated divergence caused by hypermutable methylated CpG sites [67], we excluded all the CpG sites from our calculation of LTR-LTR divergence. We applied this method to calculate the age of all complete (i.e. with two LTR) proviruses detected in cat, *M. lucifugus* and pangolin genome assemblies. We use previously estimated neutral substitution rates of 2.7×10^-9^ and 1.8×10^-9^ per year for vespertilionid bats and felids respectively [43,68], and an “average” mammal neutral substitution rate of 2.2×10^-9^ per year [69] for the pangolin.

The results of these calculations predict that the oldest MLERV1 and FcERV_γ6 proviruses would be ∼10 MY and ∼20 MY respectively (Fig. 3a). The amplification of the bat MLERV1 family would have peaked sharply in the last 2 MY, while the cat FcERV_γ6 elements inserted more continuously over the past ∼15 MY (Fig. 3a). The two MPERV1 proviruses identified in the pangolin genome are estimated to be ∼10 and ∼18 MY based on this approach.

**Fig. 3. Dating individual proviral insertions based on LTR-LTR divergence.**

(a) Age distribution of proviral insertions inferred from LTR-LTR divergence. The *y* axis shows the number of insertions for each age class binned in MY on the *x* axis. Each ERV family is shows as bars of different colors. (b) Evidence of ‘gene’ conversion between 5’ and 3’ LTR of the same provirus. Four LTR trees are shown for four pairs of orthologous proviruses shared by *M. lucifugus* (MLERV) and *M. brandtii* (MBERV). Each maximum likelihood tree was built from a multiple alignment of the 5’ and 3’ LTRs from each provirus rooted with a non-orthologous LTR from *M. lucifugus* (not shown here to simplify the figure). The ML posterior probability is shown for each node. The fact that 5’ and 3’ LTR from the same provirus tend to group together rather than by species is indicative of gene conversion between the LTRs along the two species lineages following proviral insertion in their common ancestor.

While these estimates are consistent with independent ERV invasions of the vespertilionid, felid and pangolin lineages, we noticed that the age of individual insertions based on LTR divergence were generally lower than those estimated based on their presence/absence at orthologous position across species. For instance, we found that 27 FcERV_γ6 proviruses were orthologous in cat and tiger, which indicates that all must have inserted prior to speciation of these felids, which has been robustly estimated at ∼10.8 (8.4-14.5) MY [60]. However only 13 of these 27 insertions were estimated to be older than 10 MY based on LTR divergence (S1 Table). Similarly, *M. lucifugus* and *M. brandtii* are thought to have diverged ∼10 MYA [61,63], but the age of the four MLERV1 insertions orthologous between these two species was estimated to be 10.5, 6.8, 4.5 and 1.2 MY based on LTR divergence (S1 Table). One possible explanation for these discrepancies between the two dating methods is the phenomenon of gene conversion between two adjacent LTRs, which homogenizes their sequence, causing to underestimate the date of proviral insertion [70,71]. Indeed, a phylogenetic analysis of the LTR sequences from the four MLERV1 proviruses orthologous in *M. lucifugus* and *M. brandtii*, shows a topology consistent with gene conversion for at least two of these proviruses: their 5’ and 3’ LTR cluster together rather than by species (Fig. 3b). Thus, estimates of the age of proviruses based on LTR divergence should be interpreted with caution, as they are likely to be underestimates. Nonetheless, the results are in agreement with the other lines of evidence that the vespertilionid, felid, and pangolin lineages were independently infiltrated by the same ERV during an evolutionary timeframe ranging from ∼25 to ∼13 MYA.

### Phylogenetic analysis of FcERV_γ6, MLERV1 and MPERV1 families

To further characterize the evolution history of MLERV1, FcERV_γ6 and MPERV1, we examined the phylogenetic relationship of elements within these families using a maximum-likelihood tree built from an alignment of their 3’ LTR sequences. We used only ‘complete’ proviruses (30 in bats, 43 in cats and 2 in pangolin), since we observed that including tiger provirus and solo LTRs did not yield any new major clade in the phylogeny (S1 Fig.). Also the general topology of the tree is identical if the 5’ LTR sequences are included (S2 Fig.). Trees generated using internal coding sequences also displayed the same general topology (S3 Fig.), but offered less phylogenetic resolution due to the more constrained nature of retroviral coding sequences relative to LTRs [72,73].

The unrooted tree resulting from the phylogenetic analysis (Fig. 4) clearly shows that FcERV_γ6 and MPERV1 elements are more closely related to each other than to the bat MLERV1 elements. Another striking observation is that elements within the FcERV_γ6 family fall within a single clade with uniformly short branches, whereas the MLERV1 elements, as we previously reported [41] can be divided into 3 distinct subfamilies separated by long branches, with MLERV1_2 and MLERV1_3 being closer to each other and more distant from FcERV_γ6 than MLERV1_1 (Fig. 4). These data is consistent with a scenario whereby the FcERV_γ6 family was amplified from a single infectious progenitor, while MLERV1 elements might have originated from at least three distinct infectious progenitors.

**Fig. 4. Phylogenetic analysis of MLERV1, FcERV_γ6, MPERV1 families.**

A maximum likelihood phylogenetic tree built from a multiple alignment of 3’ LTR sequences of 75 proviruses. The ML posterior probabilities supporting clades discussed in the text are shown. Information on the species origin as well as the presence/absence of envelope sequence is labeled at each node. Two independent losses of envelope by deletion are highlighted.

### Selection analysis on coding sequences reveal different amplification dynamics

To further explore the history of the FcERV_γ6 and MLERV1 families, we next turn to an analysis of selection regimes that have acted on their coding sequences during their amplification. Such analysis can help discern whether ERVs have spread primarily through reinfection or retrotransposition events because the latter mechanism, which is strictly intracellular, is predicted to be associated with relaxed constraint on coding domains normally required for infection, such as the envelope and Gag matrix domains. Indeed, the Gag matrix proteins target the plasma membrane to facilitate the budding of infectious viral particles [74,75], while the envelope protein binds to membrane receptor to promote virion entry in the host cell [5]. Thus, proviruses that originate from infection events should show evidence of functional constraint on both Gag matrix and envelope domains [76,77]. To perform this analysis, we used all MLERV1 (n=30) and cat FcERV_γ6 (n=43) proviruses with complete (or nearly complete) coding capacity. Given that only 2 proviral MPERV1 copies could be identified, we did not perform selection analysis for this family.

To evaluate how natural selection may have constrained the different coding regions of MLERV1 and FcERV_γ6, we computed the dN/dS ratio (ω) applying the branch model implemented in PAML, where dN denotes the non-synonymous substitution rate and dS denotes the synonymous substitution rate, along the branches of the phylogeny of FcERV_γ6 and MLERV1 elements for each of their predicted coding domains [78]. ω values significantly smaller than 1 are indicative of purifying selection acting to maintain a functional protein sequence, while ω values not significantly different from 1 are indicative of neutral evolution or relaxed functional constraint. To test for significant deviation of ω from 1, we apply a likelihood ratio test [79].

Within the FcERV_γ6 family, the analysis reveals that purifying selection has acted on all coding domains (ω value ranging from ∼0.6 to ∼0.9, *p* < 0.05), with the notable exception of Gag matrix and envelope domains (Fig. 5a). The ω value is not significantly different from 1 (neutral evolution) for the Gag matrix domain (Gag_MA). Besides, all but nine of the 43 FcERV_γ6 proviruses lack an envelope domain (TLV_coat). The nine copies that have retained a recognizable envelope domain occupy basal branches in the phylogeny (Fig. 4) and have orthologs in the tiger genome suggesting that they predate the envelope-less copies (S1 Table). Furthermore, 29 or the 43 FcERV_γ6 proviruses examined, including all cat-specific copies, share the same deletion breakpoint removing most of envelope gene (Fig. 5b). These data suggest that FcERV_γ6 copies potentially coding an envelope were inserted prior to the speciation of cat and tiger (∼10.8 MYA), while copies integrated more recently lacked the envelope domain. In addition, the envelope open reading frames of these nine ancient FcERV_γ6 elements accumulated multiple indels or missense substitutions. Thus, none of the FcERV_γ6 elements in the cat genome appear to have retained a functional envelope domain. Together with the lack of purifying selection acting on the Gag matrix domain, these data suggest that FcERV_γ6 rapidly lost its infectious capacity in the cat lineage but has continued to amplify primarily via retrotransposition amplified primarily via retrotransposition.

**Fig. 5. Selection analysis on coding domains**

(a) dN/dS ratio (ω) of each coding domain in FcERV_γ6, MLERV1_2 and MLERV1_3. MA, CA, PRO, RT, RH, INT and TM denote matrix, capsid, aspartyl protease, reverse transcriptase, RnaseH, integrase and envelope transmembrane domain, respectively. Asterisks denote the level of significance of departure from ω =1 (likelihood ratio test, see Methods) with * = p< 0.05; ** = p< 0.01; *** = p< 0.001. NS= not significant (p>0.05); NA = not applicable (domain deleted). (b) Shared breakpoints at the site of envelope deletion in a subset of FcERV_γ6 elements. A schematic of the prototypical proviral coding regions showing the approximate position of the envelope deletion in 29 FcERV_γ6 elements marked with blue triangles in Fig. 4 and (below) an alignment with a subset of envelope-containing FcERV_γ6 elements, showing that they share the same deletion breakpoints. These data indicate that these 29 elements likely arose from amplification of a progenitor copy that had suffered a large deletion in the envelope region.

**Fig. 5.**
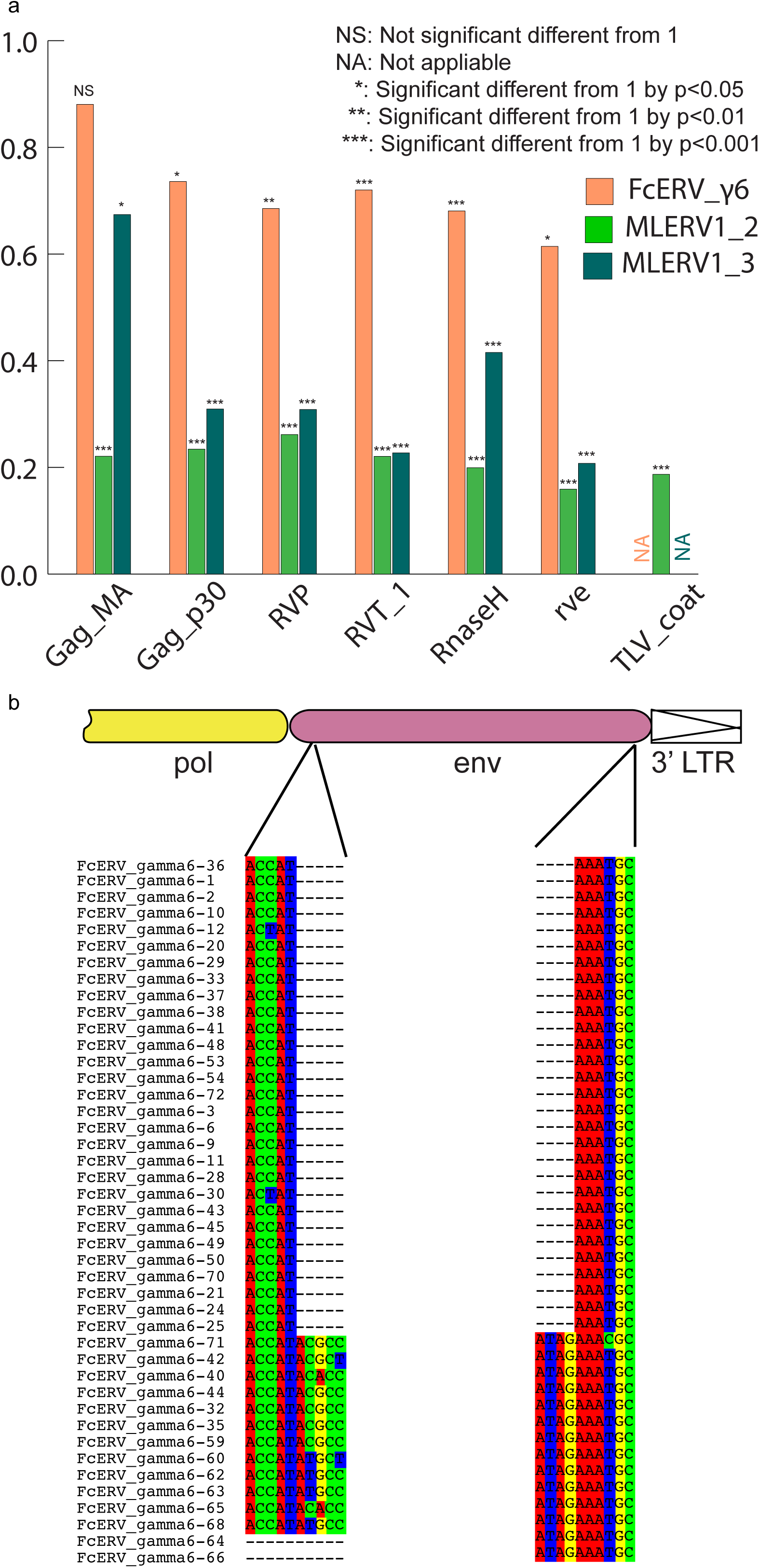
Selection analysis on coding domains. a. dN/dS ratio (ω) of each coding domain in FcERV_γ6, MLERV1_2 and MLERV1_3. MA, CA, PRO, RT, RH, INT and TM denote matrix, capsid, aspartyl protease, reverse transcriptase, RnaseH, integrase and envelope transmembrane domain, respectively. Asterisks denote the level of significance of departure from ω =1 (likelihood ratio test, see Methods) with * = p< 0.05; ** = p< 0.01; *** = p< 0.001. NS= not significant (p>0.05); NA = not applicable (domain deleted). b. Shared breakpoints at the site of envelope deletion in a subset of FcERV_γ6 elements. A schematic of the prototypical proviral coding regions showing the approximate position of the envelope deletion in 29 FcERV_γ6 elements marked with blue triangles in Fig. 4 and (below) an alignment with a subset of envelope-containing FcERV_γ6 elements, showing that they share the same deletion breakpoints. These data indicate that these 29 elements likely arose from amplification of a progenitor copy that had suffered a large deletion in the envelope region.

By contrast, selection analysis suggests that the MLERV1 family has experienced a more complex amplification history. We focused our analysis on the MLERV1_2 and MLERV1_3 subfamilies because they are the two best-supported monophyletic subfamilies with sufficient number of proviruses to draw solid conclusions. First, we observe that generally the signature of purifying selection is more pronounced on the bat elements than on the cat elements, as indicated by much lower ω values (Fig. 5a). The only exception is the Gag matrix domain of the MLERV1_3 subfamily, which exhibits relatively higher ω value (ω=0.67, p=0.02) indicative of weaker purifying selection (Fig. 5a). In addition all MLERV1_3 elements appear to have lost their envelope domain through the same deletion event (Fig. 4). This pattern contrasts with elements within the MLERV1_2 subfamily, for which all coding domains, including Gag matrix and envelope, have evolved under strong purifying selection during the spread of these elements (ω from 0.15 to 0.27, p< 0.001) (Fig. 5a). These data suggest a scenario whereby MLERV1_3 has amplified primarily by retrotransposition, while the spread of MLERV1_2 has been driven by multiple infection events (reinfection).

## DISCUSSION

### Cross-ordinal transmission of a mammalian retrovirus

Until recently, most retroviral CST events that have been documented rigorously have implicated closely related species [6,9], suggesting that the phylogenetic distance between species is an important determinant of the host range of a retrovirus [7,29]. Indeed, many previous studies have illustrated how the divergence of host cellular factors that either facilitate or restrict viral replication can modulate the host range of a virus [80-82]. The systematic analysis of retroviruses fossilized in the genome as ERVs is progressively revealing a more nuanced picture whereby some retroviruses appear to have been capable to infect widely diverged species (i.e. belonging to different orders) without seemingly much changes occurring in their own sequences. Recent large-scale pylogenomics analyses have suggested that cross-order transmission may actually be fairly common for some groups of retroviruses, and in particular gammaretroviruses [19,20,33]. While these studies disclosed phylogenetic patterns suggestive of CST events, they did not explicitly rule out alternative hypotheses, such as vertical persistence and stochastic loss of the ERV in some lineages, and thus they generally await confirmation through more detail analyses such as the one presented here.

Our study provides multiple lines of evidence supporting the notion that a gammaretrovirus infiltrated independently the germline of bat, cat and pangolin species representing three mammalian orders (Chiroptera, Carnivora, Pholidota, respectively). First, elements found in these species display a level of nucleotide sequence similarity (∼85%) along their entire length that is comparable to that observed between closely related retroviruses that have undergone very recent CST (such as SIVcpz and HIV-1). Such a level of sequence similarity between ERVs inhabiting species diverged by ∼85 MYA [53] is incompatible with a scenario of vertical descent from an ERV inherited from their common ancestor. The CST hypothesis is also bolstered by the highly discontinuous taxonomic distribution of this particular ERV family. Out of 107 mammal species for which whole genome assemblies are publicly available, we could only detect members of this ERV family in vespertilionid bats, felids and pangolin, but not in several species representing related mammal families (6 additional Chiroptera species from 4 families and 7 additional Carnivora species from 5 families). Thus, a scenario evoking a single introduction of this ERV family in the common ancestor of bats, cats and pangolins followed by vertical inheritance would necessitate at least 5 independent losses (Fig. 1b) to account for its current taxonomic distribution. A more parsimonious scenario is that this ERV family was acquired horizontally and independently in each of the three species lineages where it is currently detected. Finally, our estimation of the dates at which these elements first entered their host genomes, which relies on two independent approaches (cross-species comparison of orthologous ERV loci and LTR-LTR divergence), converges to a bracket of 13 to 25 MYA, which far postdates the divergence of their host species (∼85 MYA). Together these data indicate that a progenitor gammaretrovirus infiltrated, within a relatively short timeframe, the germlines of ancestral vespertilionid bat, felid and pangolin species.

It is conceivable that this retrovirus could have transferred directly between these ancestral species because their geographic distribution likely overlapped in Eurasia during the estimated period of initial ERV infiltration (∼13-25 MYA) [60,61,83]. Given that cats are known to prey on both bats [84-87] and pangolins [88], a direct transfer from bat or pangolin to cat is plausible. Indeed, predation has been put forward as the most likely explanation for the spillover of bat lyssaviruses (rabies) into domestic cats [89]. On the other end, both bats and pangolins are capable of surviving a cat attack, which makes the transfer from predator to prey conceivable as well. Nonetheless, multiple lines of evidence indicate that MLERV1 colonized these bat genomes more recently (Fig. 2 and Fig. 3), which may suggest a CST from cat to bat. Furthermore, we cannot rule out that one or several intermediate hosts were involved in the introduction of the retrovirus in these species.

### Repeated transition from retrovirus to retrotransposon

Our data suggest that, shortly after infiltration of the felid genome, FcERV_γ6 lost the capacity to infect cells and transformed into a retrotransposon. Two lines of evidence support this transition. First, the Gag matrix domain of FcERV_γ6 has been neutrally evolving while these elements spread in the cat genome (Fig. 5). The Gag matrix protein directs virus-like particles to the cellular membrane and promote retroviral particle assembly and budding [74,75,90]. Second, all the FcERV_γ6 elements amplified in the domestic cat lineage clearly derive from a progenitor that lacked coding capacity for a functional envelope protein (Fig. 4 and Fig. 5b). Together these data suggest that FcERV_γ6 lost its infectious capability soon after it became endogenous, but continued to propagate by retrotransposition, much like the IAP elements in the mouse genome [91,92]. A recent study showed that a FcERV_γ6 insertion in the *KIT* gene currently segregating in domestic cats is responsible for the “Dominant White” and white spotting pigmentation phenotypes [93], which supports our findings that some FcERV_γ6 insertion activity is very recent and likely ongoing. Interestingly, the FcERV_γ6 element inserted at the *KIT* locus lacks envelope domain and clusters with other recently active FcERV_γ6 copies in our phylogenetic analysis (Fig. 4). Collectively these data suggest that FcERV_γ6 has morphed into a successful retrotransposon that may still be active in the domestic cat.

In contrast to FcERV_γ6, the sequence diversity and phylogenetic structure of MLERV1 elements in the vesper bat genomes are indicative of a more complex amplification history characterized by a mixture of retrotransposition and reinfection events. Our phylogenetic analysis delineates a least three highly diverged MLERV1 subfamilies. Furthermore, selection analyses suggest that different subfamilies have adopted different evolutionary trajectories. The MLERV1_2 subfamily is characterized by a signature of intense purifying selection acting on all coding regions throughout the whole clade (Fig. 5a). These data strongly suggest that elements within that subfamily have retained their infectious capacities for extended period of time and most likely spread primarily through reinfection events. It is even possible that MLERV1_2 is still active and infectious: most insertions are very recent (Fig. 2 and Supplymentary Table) and at least one copy (MLERV1.80) contains apparently full-length and intact *gag*, *pol*, and *env* genes.

The MLERV1_3 subfamily appears to have followed a different evolutionary path whereby the divergence of the elements was accompanied by a strong signature of purifying selection in all coding regions with the notable exception of the Gag matrix domain where the signature of selection is much weaker (Fig. 5a) and the envelope domain which was apparently deleted altogether. This selection pattern resembles that of FcERV_γ6 and is indicative of proliferation primarily via retrotransposition as opposed to reinfection. Consistent with this hypothesis and the so-called superspreader model [94], the MLERV1_3 family has been by far the most successful at spreading during *Myotis* evolution: it has the highest copy number, including many species-specific insertions (Table 1).

Interestingly, none of the 29 MLERV1 elements identified in the big brown bat *E. fuscus* belong to the MLERV1_3 subfamily. This is consistent with the idea that the MLERV1_3 subfamily originated after the split of Eptesicus-Myotis split ∼25 MYA and amplified during the diversification of the *Myotis* lineage. At present it remains unclear whether the MLERV1 elements present in *E. fuscus* and *Myotis* descend from element(s) introduced in their common ancestor or if they result from independent acquisition of the same retrovirus. On the one hand, the observation that both *E. fuscus* and *Myotis* harbor elements from two diverged subfamilies may be interpreted as evidence that these subfamilies descend from a single progenitor ERV acquired in the common ancestor of these species. On the other hand, the fact that none of the MLERV1 insertions are shared (orthologous) between *E. fuscus* and any of the 3 *Myotis* genomes (Fig. 2) and that none of the provirus insertions dated in any of these bat species appear older than 13 MY (considerably less than the estimated divergence between the two genera, 25 MYA) (Fig. 3) supports a scenario of multiple, independent acquisition. This scenario, while requiring at least two CST events, is conceivable because *Eptesicus* and *Myotis* bats likely occupied a widely overlapping geographic distribution at the estimated time of MLERV1 invasions [61] and these congeners are currently known to frequently come into contact within the same roost [95,96].

Regardless of the origin of MLERV1, the data summarized above illustrate how the same retrovirus has infiltrated widely diverged mammals and transitioned multiple times (at least twice: FcERV_γ6 and MLERV1_3) from an infectious pathogen to a genomic parasite (i.e. a retrotransposon). The biological factors and sequence of events underlying such transition remain poorly understood. In a seminal study, Ribet et al. showed that the loss of envelope gene combined to the gain of an endoplasmic reticulum targeting signal were apparently sufficient for an infectious progenitor of the mouse IAP elements to turn into a highly active retrotransposon [90]. Magiorkinis et al. [94] have extended this paradigm and proposed that the passive loss of envelope lead ERVs to become “superspreaders” in the genome. Through a study of IAP-like elements across a wide range of species, these authors observed that envelope-less elements generally achieve much higher copy numbers than those maintaining a functional envelope. Our results support this model. First, envelope-less FcERV_γ6 elements have proliferated to high copy numbers in the domestic cat (n=832) and tiger (n=730). In addition, in the bats the only subfamily of MLERV1 elements that has attained similarly high copy number is MLERV1_3, which conspicuously lack a functional envelope gene (Fig. 5). MLERV1_3 elements have generated many species-specific insertions consistently outnumbering the MLERV1_2 subfamily, which appears to have spread primarily by reinfection (659 vs. 14 in *M. lucifugus* genome, 350 vs. 12 in *M. brandtii* and 331 vs. 19 in *M. davidii*) (Fig. 2). Thus, there is a clear association between the loss of infectious capacity and ERV expansion by retrotransposition.

### Does host biology affect ERV proliferation?

An intriguing finding of this study is that the same or very similar retrovirus was endogenized in three different mammalian hosts within a relatively short evolutionary timeframe (∼15-25 MYA), but followed quite different evolutionary trajectories in the three species lineages. In the pangolin lineage, the ERV family failed to amplify (only 2 detectable copies) and was essentially ‘dead-on-arrival’. In the cat lineage, the ERV progenitor apparently lost its infectious capacity shortly after endogenization and subsequently amplified to high copy numbers by retrotransposition through an extended period of time ranging from at least 10 MYA (256 insertions orthologous in cat and tiger) to modern times (*KIT* insertion segregating in domestic cats). Meanwhile, in the bat lineage, the ERV followed a more complex evolutionary path characterized by multiple episodes of reinfection, and at least one burst of amplification by retrotransposition. These observations beg the question whether the loss of infectious capacity of an ERV and its conversion to a retrotransposon is a purely stochastic process, largely owing to the stochastic loss of envelope and gag matrix functions, or if it can be influenced by some biological characteristics of the host species? The pattern of sustained reinfection of MLERV1 in the bat lineage is particularly intriguing in light of the growing awareness that bats seem to frequently act as reservoir for viruses otherwise lethal to other mammals [34,38]. The reasons for bats’ propensity to support high and diverse loads of viral pathogens are poorly understood, but it is thought that some physiological (e.g. immunopathological tolerance) and/or ecological features (e.g. flight, roosting) allow these animals to tolerate higher level of viral replication and/or facilitate viral transmission [38,97,98]. By the same token, it is tempting to speculate that the same properties might predispose bats to support higher level of ERV reinfection compared to other mammals such as cats. Testing this hypothesis will necessitate a more systematic examination of the amplification dynamics of ERVs in a wide range of mammals to assess whether the tendency toward maintenance of infectious capacity is a general trademark of bats or possibly other groups of mammals.

## METHODS

### Initial detection of CST events involving MLERVs

Nucleotide sequences of all RVT_1 domains of previously identified MLERVs [1,41] were used as queries to search the whole genome sequence database from the National Center for Biotechnology Information (NCBI) using default MegaBLAST parameters [99]. An 80% similarity over 80% region was used as filter to exclude non-specific hits.

### Identification of complete proviruses, putative full-length ERVs and solo LTRs

Complete MLERV1 and FcERV_γ6 proviruses in the *M. lucifugus* and cat genomes were collected from previous publications [41,48]. To ensure we only considered elements from these families, we only retained elements with 80% nucleotide similarity to another family member, a procedure which resulted in the exclusion of the FcERV_γ6_46 copy from the FcERV_γ6 family (S3 Fig.).

To identify complete proviruses in other vesper bat genomes, the RVT_1 domain sequence of MLERV1.71 in *M. lucifugus* was used as query in blastn search of the *M. brandtii*, *M. davidii* and *E. fuscus* genome assemblies available in NCBI. In parallel, we applied LTRharvest [100] and LTRdigest [101] as described previously [41] to identify all putative proviruses in each of the three bat genome assemblies. We then used BEDTools to intersect the coordinates of RVT_1 domain blastn hits with that of the candidate proviruses [102]. All the candidate proviruses intersecting with a MLERV1 RVT_1 hit were ‘manually’ inspected to refine their termini and confirm their identity as members of the MLERV1 family.

To comprehensively retrieve all proviruses and solo LTRs related to the FcERV_γ6/MLERV1/MPERV1 families in each of their respective genomes, we run RepeatMasker [103] with default setting and a custom repeat library with representitives from all MLERV1/MPERV1/FcERV_γ6 subfamilies against each genome assembly. The RepeatMasker output was then parsed using script parse_RMout_count_solo_and_full.pl to produce bed files of all complete solo LTRs and full length ERVs. We define a complete solo LTR as a sequence matching the LTR with missing less than 150 bp at their 5’ termini and missing less than 10 bp at their 3’ termini. We identified elements as putative proviruses those delimited by two LTRs in the same orientation separated by 3 kb to 10 kb of intervening sequence. Manual inspection of a subset of putative proviruses identified by this approach confirmed that most contained typical ERV coding sequences, though frequently interrupted by large sequence/assembly gaps. The LTR libraries and PERL scripts used for these analyses have been deposited on Github (https://github.com/xzhuo/orthologusLTR.git).

In the pangolin genome, our initial MEGAblast search yielded only 2 significant hits to the MLERV1 RVT_1 domain, but many more related ERVs could be retrieved using the RT domain from these two initial hits in reiterative blast searches against the pangolin genome assembly. To examine the relationship of these RT elements to each other and to the MLERV1 and FcERV_γ6 families, we conducted a phylogenetic analysis using the Maximum Likelihood package PhyML3.1 with the GTR+Γ model [104]. The resulting tree (S3 Fig.) revealed that only the two initial hits clustered with FcERV_γ6/MLERV1 and were considered part of the MPERV1 family. The other elements form a distinct family we called MPERV2. MPERV1 and MPERV2 elements share less than 80% nucleotide sequence similarity in their coding regions, but still retain substantial level of sequence similarity in their LTRs. Thus, to correctly estimate the number of solo LTRs for the MPERV1 family, we had to examine their position on a phylogenetic tree of LTRs (data not shown). Using this approach, 27 solo LTRs could be assigned to the MPERV1 family in the pangolin genome assembly (as reported in Table 1). Because MPERV2 was much more distantly related to FcERV_γ6 and MLERV1 (< 80% sequence similarity), we did not analyze further MPERV2 in this study. Reference sequences for MPERV1 and MPERV2 have been deposited in Repbase [47].

All identified ERVs are available as bed format (S1 File).

### Sliding window pairwise similarity calculation

To generate the sliding window analysis shown in Fig. 1a, we used MUSCLE [105] to align the nucleotide sequence for three pairs of proviruses: MLERV1_77 vs. FcERV_γ6_62; FcERV_γ6_62 v.s MPERV1_ltr106; SIVcpz(AF115393) vs. HIV-1(NC_001802) and used SEAVIEW [106] to manually adjust each alignment. Each pairwise alignment was then split into non-overlapping windows of 50 bp (i.e. =175 segments for the MLERV1_77 vs. FcERV_γ6_62 alignment) and the percentage of sequence similarity was computed for each window.

### ERV orthologus loci identification

Orthologous ERV loci were detected similarly as we described previously [41]. Briefly, we used the Perl script extract_flanking_fasta.pl to extract 200 bp at both ends of each query element along with 200 bp of flanking sequences. The output file is then used as query in a batch blastn search against the target genome assembly with default parameters. The csv format blast output is then parsed using orthoblast_finder.pl to pair 5’ end hits with 3’end hits. Finally, the paired hits output was parsed using the script final_annotation.pl to infer the presence/absence of each element in the target genome. All these perl scripts were deposited on Github (https://github.com/xzhuo/orthologusLTR.git).

### Estimation of individual provirus insertions using LTR–LTR divergence

Sequence divergence between 5’ and 3’ LTRs from the same provirus was computed as previously described [41]. To infer insertion dates from LTR divergence of MLERV1 and FcERV_γ6 elements, we used previously estimated lineage-specific neutral substitution rates of 2.7 × 10^-9^ yr^-1^ [68] and 1.8 × 10^-9^ yr^-1^ [43] for the vespertilionid and felid lineages respectively. Since no substitution rate has yet been estimated for the pangolin lineage, we used the ‘average’ mammal neutral substitution rate of 2.2 × 10^-9^ yr^-1^ [69] to infer the age of MPERV1 insertions.

### Phylogenetic analysis

The maximum-likelihood phylogenies presented for LTR sequences were built using PhyML3.1 [107]. The multiple alignment of LTR sequences was constructed using MUSCLE with default nucleotide parameters and manually adjusted using SEAVIEW [105,106]. Nucleotide substitution model was chosen using AIC criterion in jmodeltest2.1.6 [108] (GTR + Γ). Dendroscope 3 was used for tree visualization [109].

### Selection analysis on coding domains

ERV coding regions were predicted using HMMER3 in all 6 reading frames [110] to delineate Gag_MA (matrix), Gag_p30 (capsid), RVP (protease), RVT_1 (reverse transcriptase), RnaseH, rve (integrase) and TLV_coat (envelope transmembrane domain) domains in MLERV1, FcERV_γ6 and MPERV1 proviruses. A multiple codon alignment was generated for each set of coding domains using MUSCLE and manually adjusted with SEAVIEW [106]. The program codeml from the PAML4.8 package [78] was used to estimate dN/dS ratio with branch model = 2. A maximum likelihood phylogeny of the LTR sequences was used as the guide tree in codeml. To test for purifying selection on each coding domain, we calculated the control lnL value by running codeml with ω fixed to 1. Then likelihood ratio test was performed as suggested by PAML to test if ω is significantly different from 1 [79].

## ACKNOWLEDGEMENTS

We thank Aurelie Kapusta, Ray Malfavon-Borja, Edward Chuong, John McCormick, Ellen Pritham, Claudia Marquez and Rachel L. Cosby for helpful discussion.

